# The expanded BXD family of mice: A cohort for experimental systems genetics and precision medicine

**DOI:** 10.1101/672097

**Authors:** David G. Ashbrook, Danny Arends, Pjotr Prins, Megan K. Mulligan, Suheeta Roy, Evan G. Williams, Cathleen M. Lutz, Alicia Valenzuela, Casey J. Bohl, Jesse F. Ingels, Melinda S. McCarty, Arthur G. Centeno, Reinmar Hager, Johan Auwerx, Saunak Sen, Lu Lu, Robert W. Williams

**Author notes:** Contributed equally to this work, and should both be considered as joint first authors (David G. Ashbrook, Danny Arends). Corresponding authors: David G. Ashbrook,; Robert W. Williams,; Lu Lu. Author email & ORCID David G. Ashbrook -, Danny Arends -, Pjotr Prins -, Megan K. Mulligan -, Suheeta Roy -, Evan G. Williams -, Cathleen M. Lutz -, Alicia Valenzuela -, Casey J. Bohl -, Jesse F. Ingels -, Melinda McCarty -, Arthur G. Centeno -, Reinmar Hager -, Johan Auwerx -, Saunak Sen - - 0000-0003-4519-6361, Lu Lu -, Robert W. Williams.

## Abstract

The challenge of precision medicine is to model complex interactions among DNA variants, sets of phenotypes, and complex environmental factors and confounders. We have expanded the BXD family, creating a powerful and extensible test bed for experimental precision medicine and an ideal cohort to study gene-by-environmental interactions.

These BXD segregate for over 6 million variants, with a mean minor allele frequency close to 0.5. We have increased the family two-fold to 150 inbred strains, all derived from C57BL/6J and DBA/2J. We have also generated updated and comprehensive genotypes and an unrivaled deep phenome.

Approximately 10,000 recombinations have been located, allowing precision of quantitative trait loci mapping of ±2.0 Mb over much of the genome and ±0.5 Mb for Mendelian loci. The BXD phenome includes more than 100 ‘omics data sets and >7000 quantitative and clinical phenotypes, all of which is publicly available.

The BXD family is an enduring, collaborative, and replicable resource to test causal and mechanistic links between genomes and phenomes at many stages and under a wide variety of treatments and interventions.

## Background

### The origin of the BXD family

Recombinant inbred strains of mice, and the BXD family in particular, have been used for 45 years to map Mendelian and quantitative trait loci ^1–5^. Production was started in 1971 by Benjamin A. Taylor by crossing a female C57BL/6J (B6 or B) and a male DBA/2J (D2 or D)—hence BXD. The first set of 26 (expanded in the 1990s to 35) recombinant inbred strains were intended mainly for mapping Mendelian loci ^1, 6^, but the family was soon used to map more complex quantitative traits—cancer susceptibility ^7–9^, neuroanatomical traits ^10–13^, and behavioral differences ^14, 15^, and pharmacological responses to toxicants and drugs ^16–19^. All of these strains are still available from The Jackson Laboratory (JAX) and carry the strain suffix “/TyJ”.

Production of BXD43 through BXD102 started in the late 1990s at the University of Tennessee Health Science Center (UTHSC) ^20, 21^. These new strains were derived from advanced intercross (AI) progeny that had been bred for as many as 14 generations before inbreeding ^22^ (Figure 1; Supplementary figure 1; Supplementary table 1). AI-derived BXDs incorporate roughly twice as many fixed cross-over events (recombinations) between *B* and *D* parental genomes compared to F2-derived BXDs—80 versus 40 ^22–26^ (Figure 2). This improved both power and precision.

**Figure 1:**
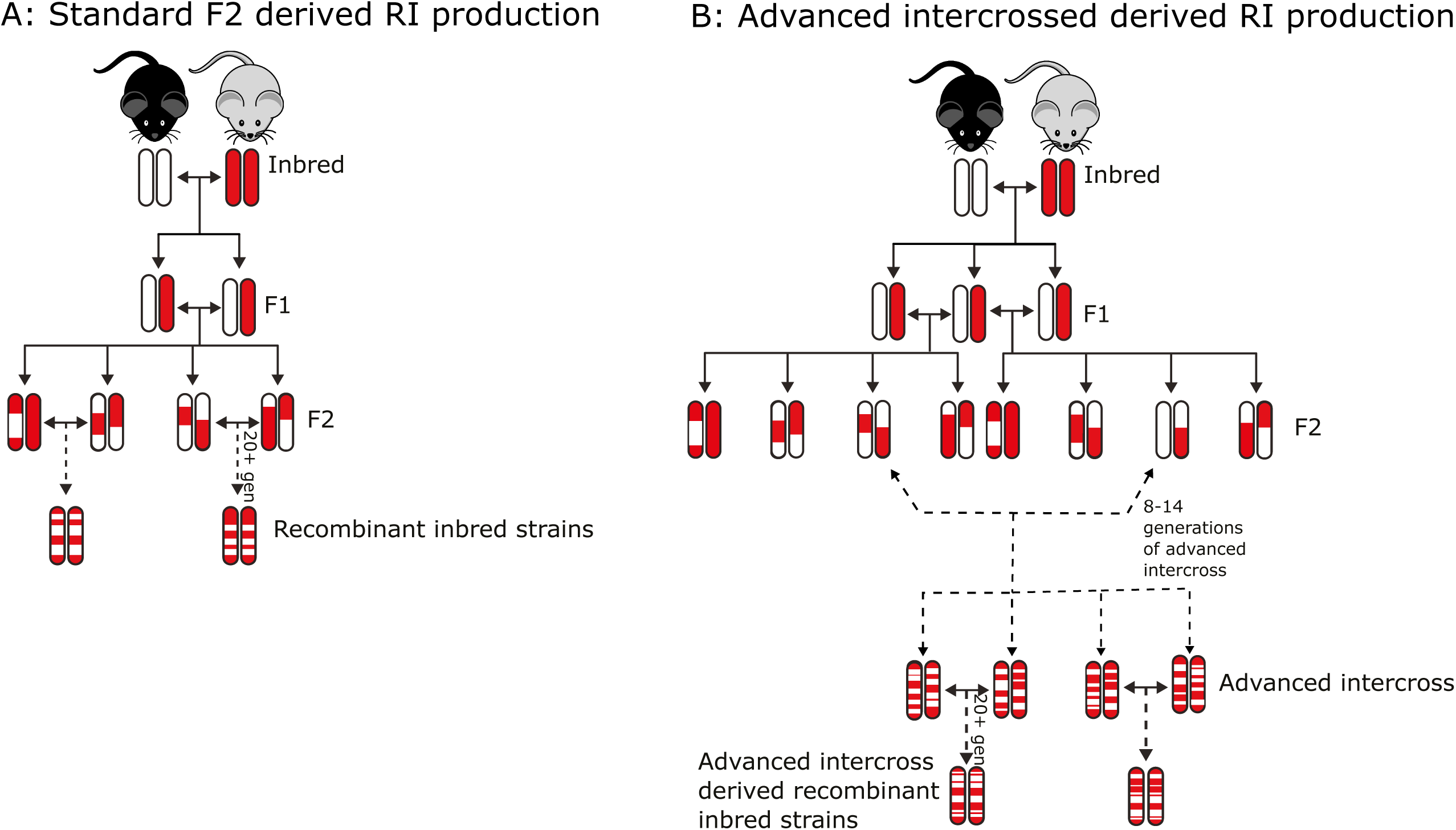
Production of the BXD recombinant inbred family by either standard F2 (A) or advanced intercross (B). Four epochs of the BXD were derived from F2 (totaling ∼75 strains) and two epochs were derived from advanced intercross (totaling ∼65 strains). Red coloring has been used to represent regions of the genome coming from the inbred C57BL/6J (B6) parental strain, whereas white coloring has been used to represent regions of the genome coming from the inbred DBA/2J (D2) strain. Solid lines have been used to represent a single generation, whereas dashed lines represent several generations, with the number of generations written along the line. Full details can be found in Supplementary table 1 and Supplementary figure 1. Adapted from (Nadeau and Auwerx, 2019; Peirce et al., 2004; Williams and Auwerx, 2015)

**Figure 2:**
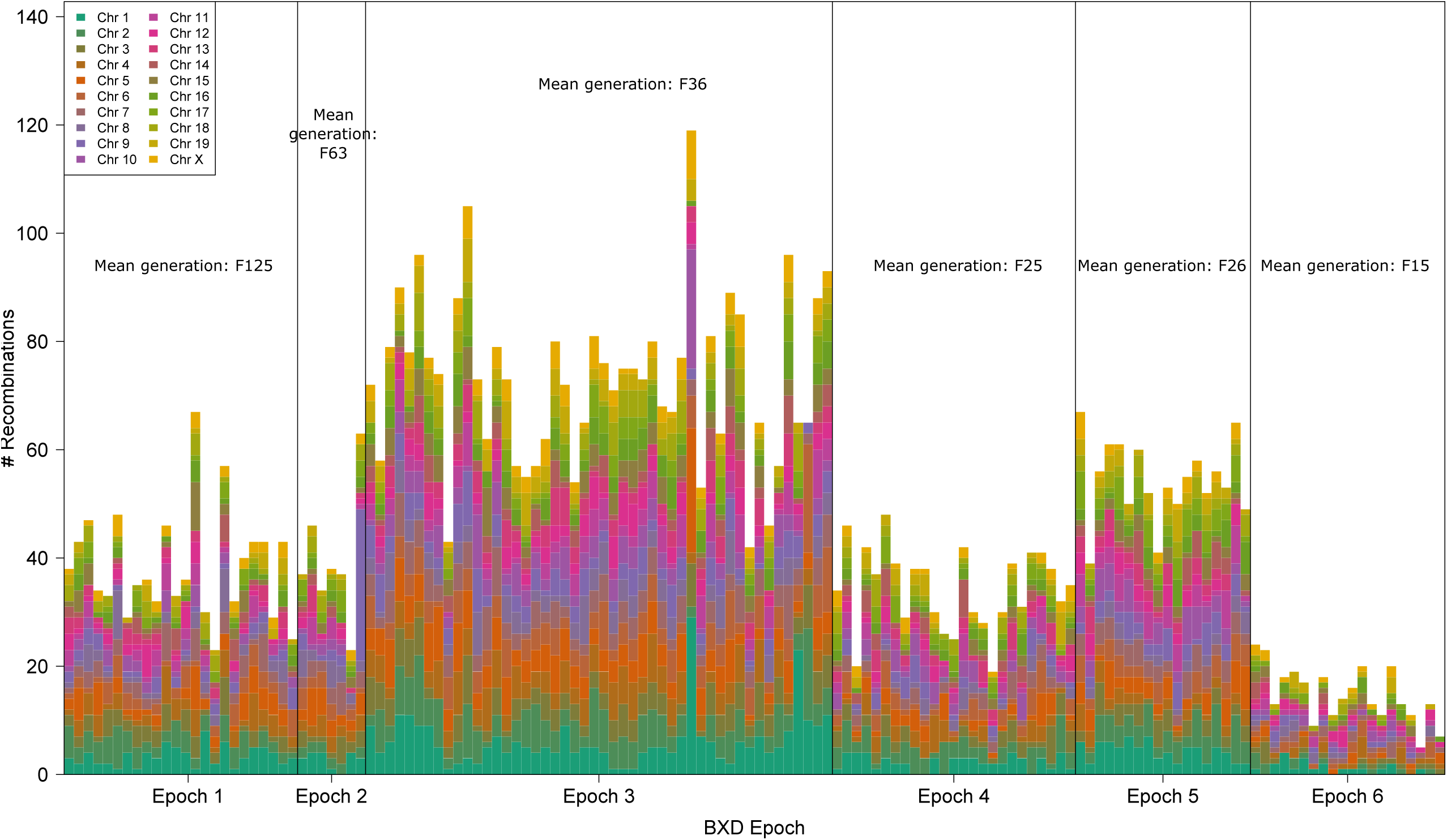
Genome wide visualization of recombinations in the BXD family. For each chromosome we computed the number of observed recombinations per strain since the start of the chromosome (nRecS), ignoring the heterozygous and unknown genotypes. The number of recombinations for each BXD strains per chromosome was plotted using a color gradient. Strain numbers are used for the ordering on the X-axis, horizontal lines separate the different epochs, and these epochs are annotated on the X-axis. The mean number of inbreeding generations is shown each epoch. Full details of the epochs can be found in Supplementary figure 1 and Supplementary table 1. Note epoch 6 appears to have fewer recombinations due to a large number of heterozygous loci at the time of genotyping.

### The expanded BXD family

Here we report on the production of new sets of BXD strains that double the family size to a total of 150 inbred stains. All have been genotyped using high-density SNP arrays. We have assembled genetic maps that improve power and mapping precision. What sets the expanded BXD family apart from all other resources for systems genetics is an unrivalled, dense and well-integrated phenome, of over 7000 classical phenotypes and over 100 ‘omics datasets. This open and publicly available phenome extends back to Taylor’s 1973 analysis of cadmium toxicity through to recent quantitative studies of metabolism ^3, 4, 27, 28^, addiction ^29–31^, behavior ^32–35^, the complex genetics of infectious disease ^36, 37^, and even indirect genetic effects ^38–40^. The BXDs have been used to test specific developmental and evolutionary hypotheses ^10, 41, 42^, to study gene-by-environmental interactions (GXE), and to quantify the consequences of interventions and treatments as a function of genome, age, and sex ^5^. We have also assembled deep and extensive ‘omics resources, including full sequence for both parents ^3, 27, 37, 43, 44^, access to almost all BXD expression and phenotype data freely available on open source web services, and powerful statistical tools designed for global analysis (e.g. GeneNetwork.org and Systems-Genetics.org) ^5, 45, 46^. Use of the expanded BXDs has accelerated. There are currently more than 500 BXD-related publications, of which more than 100 were published in the last five years.

Compared to conventional F2 and advanced intercrosses (AIs), outcrossed heterogenous stock, or diversity outbred stock, the BXD are particularly advantageous when the heritability of a trait is moderate or low because genetic signal can be boosted greatly by resampling isogenic members of the same line many times ^47^. The main benefit however is that those using BXDs gain access to coherent genomes and quantitative phenomes generated under diverse laboratory and environmental conditions ^48^. New data can be compared to thousands of publicly available quantitative traits, and with each addition, the number of network connections grows quadratically—enabling powerful multi-systems analysis for all users.

Genetic research is moving toward predictive modeling of mechanisms, health status and disease risk, and the efficacy of interventions ^49^. The expanded BXD family is extremely well suited to address more integrative questions that encompass both high levels of genetic and environmental variation. The family is also extendible into a massive diallel cross (DX) made up of 23,104 (152^2^) isogenic lines, all of which have an entirely defined genome, and any subset of which can be generated efficiently for *in vitro* and *in vivo* predictive biology and experimental precision medicine.

## Results

### The current phase of BXD production

We initiated 108 new strains between 2009 and 2013 (BXD104–BXD220). Of these, 49 are now at filial genertion 20 (F20) or higher. Inbreeding depression is a serious problem. Of 108 strains we initiated in this most recent phase of the BXD expansion, 44 are either extinct or required out crossing to save (Supplementary figure 1, Supplementary table 1). We are now testing a simple alternative in which struggling pairs of nearly inbred strains are crossed to produce more viable F1 progeny that are then used to initiate a new inbred line ^23, 50, 51^ (Supplementary figure 1).

As of March 2019, a total of 121 BXD lines are available from The Jackson Laboratory, 98 of which are listed as ‘Repository Live’ (a lead time of 0–2 weeks), 15 are listed as on pre-order (currently in production), and 8 are cryopreserved (Supplementary table 1). An additional 29 incipient BXD strains are available from the group at UTHSC, and will be donated to JAX once they are fully inbred. All BXDs are available under a standard material transfer agreement; the most important limitation being that strains cannot be sold or distributed without approval of The Jackson Laboratory or UTHSC.

### Errors during production

We have identified strains with markedly similar genotypes resulting from production errors, where a mismatched strain was crossed during production or maintainance, and these now related sets of strains have been renamed and maked by the addition of “a” and “b” suffixes. For example, BXD65, BXD65a (originally BXD97), and BXD65b (originally BXD92) are identical by descent across 90% or more of their genomes. There are important statistical consequences of non-independence and admixture of this type. For example, without corrections, the inclusion of these three BXD65 substrains gives them undue influence over results (but can be highly interesting if differences are seen between them). These production errors are different from the expected and much more subtle relations among all BXD family members that results from the inevitable pedigree relatedness (identity by descent), and by genetic drift, backcrossing, and unintended selection. Fortunately, correcting for both types of non-independence is no longer difficult using advanced QTL mapping algorithms. The Genome-wide Efficient Mixed Model Association (GEMMA) algorithm (github.com/genetics-statistics/GEMMA) has been implemented in GeneNetwork2 ^46^ and performs kinship correction during mapping.

### Improved genetic mapping

Approximately 198 BXD strains have been genotyped at over 100,000 informative markers using Affymetrix and Illumina array-based platforms ^6, 20, 52, 53^. Most markers, even those that are nominally informative, provide redundant information. Until 2016, a subset of 3,830 markers were used for mapping of phenotypes in BXD1 to BXD102. In this study we have increased the number of useful markers to 7,324. This set of markers either have unique genotype patterns (also known as “strain distribution patterns”) or define the proximal or distal limits of chromosomal intervals that are non-recombinant (plateau regions). Markers are spaced at an average of 0.63 Mb ± 1.0 SE, closely matching the asymptotic map resolution of ± 1 Mb. There are still ∼200 gaps of 2 Mb or greater without informative markers. In all cases these regions are almost identical by descent between parental strains. Five of the largest gaps are on Chr X (from 8 to 25 Mb), and other gaps are on Chrs 7, 8, 9, 14 and 18.

Moving from 3830 to 7324 markers has significantly improved the ability to detect significant linkage. For instance, Figure 3 contrast maps for *Ectromelia* virus-induced mortality (GeneNetwork trait 10043) mapped using either the older and smaller set of markers (Figure 3A) or the newer and larger set of markers (Figure 3B). A previously non-significant QTL for this trait originally published in 1991 ^54^ now achieves genome-wide significance.

**Figure 3:**
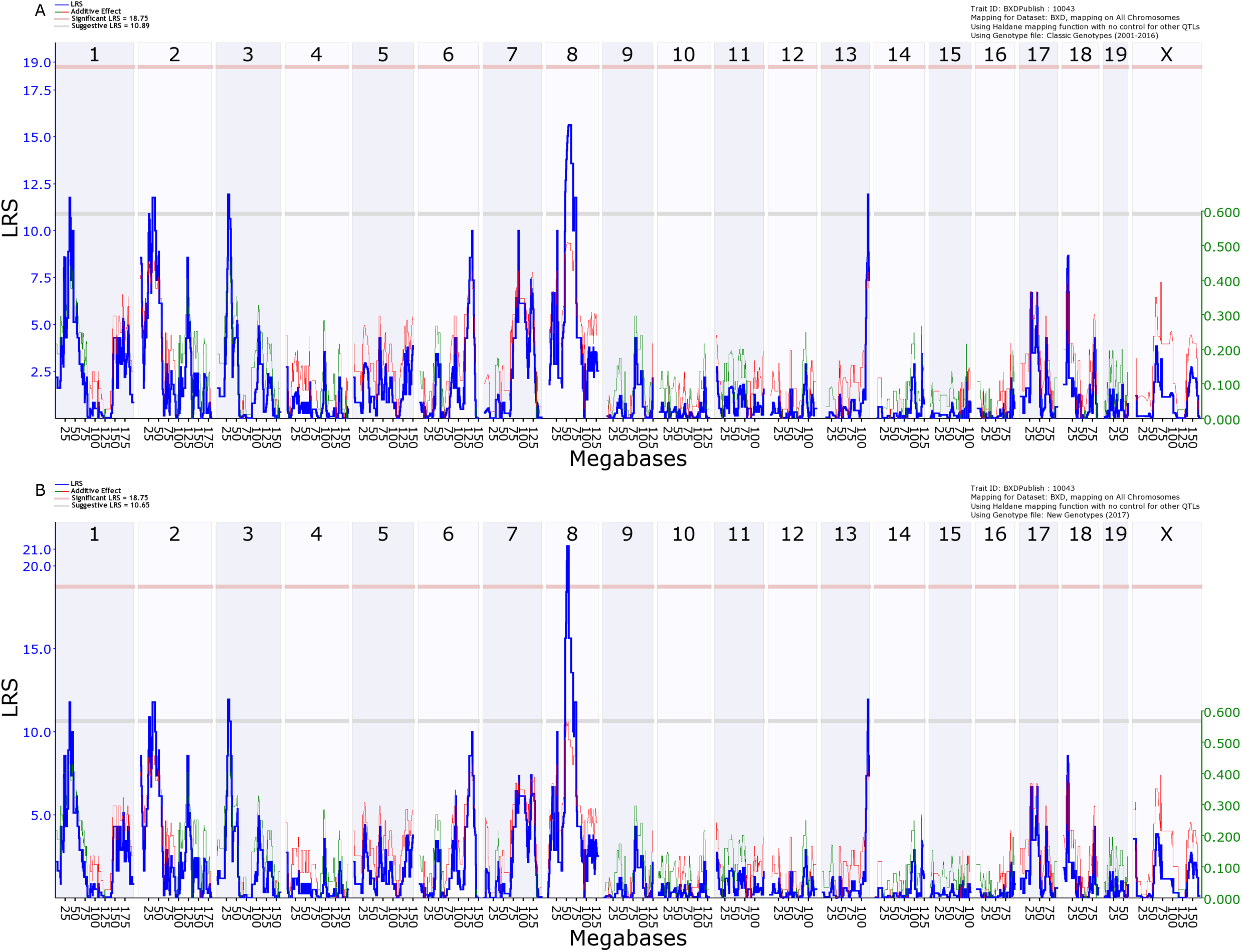
Improved mapping using updated marker map on identical data. QTL maps for GeneNetwork phenotype 10043, *Ectromelia* virus-induced mortality in BXD males, mapped using either the ‘classic’ (2001– 2016) genotypes **(A)**, or the ‘new’ (2017+) genotypes **(B)**. The blue line represents the genome scan, showing the likelihood ratio statistic (LRS) associated with each marker across the 19 autosomal and the X chromosome. The top, pink, line marks genome-wide significance, the lower, grey, line the suggestive significance threshold. The green or red lines show the additive effect coefficient, with green showing that *D* alleles increase trait values and red that *B* alleles increase trait values. The green axis on the right shows by how much the alleles increase trait values.

To quantify the improvement in mapping, we computed linkage for the entire BXD phenome (7562 phenotypes), using three different genotype files. We used the classic (2001–2016) and new genotypes (used since 2017), as well as the original BXD reference genotypes (pre-2001). Each update approximatley doubles the number of markers (from 1566, to 3830, and now 7324). We counted numbers of phenotypes with likelihood ratio statistic (LRS) scores at three cut-offs: 15 (suggestive), 20 (significant) and 25 (highly significant) (Table 1). The new genotype files increase the number of QTLs detected at suggestive and significant LRS levels, increasing the percentage detected by 17–19% and 24–26% respectively. However, there is a much smaller increase (2% and 10% respectively) at the highly significant cutoff (LRS >25) because large effect QTLs and Mendelian loci are relatively insensitive to low marker density.

**Table 1:**
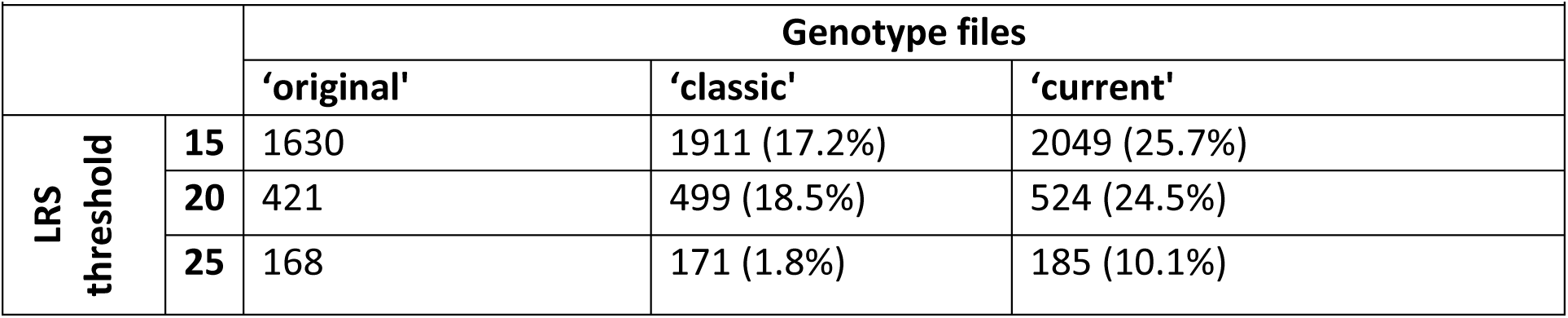
Increase in power to detect QTL using genotype files with increasing numbers of markers. For each of three genotype files with increasing numbers of markers (original = 1566 markers; classic = 3830; current = 7324) the number of phenotypes with an LRS value greater than the three threshold values (15, 20 and 25) is shown. For the classic and current genotype files, the percentage increase from the ‘original’ genotype file is also shown.

|  |  |  |  |  |
| --- | --- | --- | --- | --- |
|  |  | **Genotype files** |  |  |
|  |  | **'original'** | **'classic'** | **'current'** |
| **LRS threshold** | **15** | 1630 | 1911 (17.2%) | 2049 (25.7%) |
|  | **20** | 421 | 499 (18.5%) | 524 (24.5%) |
|  | **25** | 168 | 171 (1.8%) | 185 (10.1%) |

### Empirical mapping precision ranges from 0.25 to 5 Mb

We computed the empirical precision of mapping using a total of ∼270,000 cis-acting expression QTLs (eQTLs) with logarithm of odds (LOD ≈ –logP ≈ LRS/4.61) scores greater than 3.0 and within ±10 Mb of the cognate gene (Figure 4). This definition of a *cis*-acting eQTL is conservative in the sense that our 20 Mb acceptance window will degrade empiral estimates of precision because hits will include some regional *trans*-acting eQTLs. However, we view this approach as necessary to guard against overly optimistic estimates of precision.

**Figure 4:**
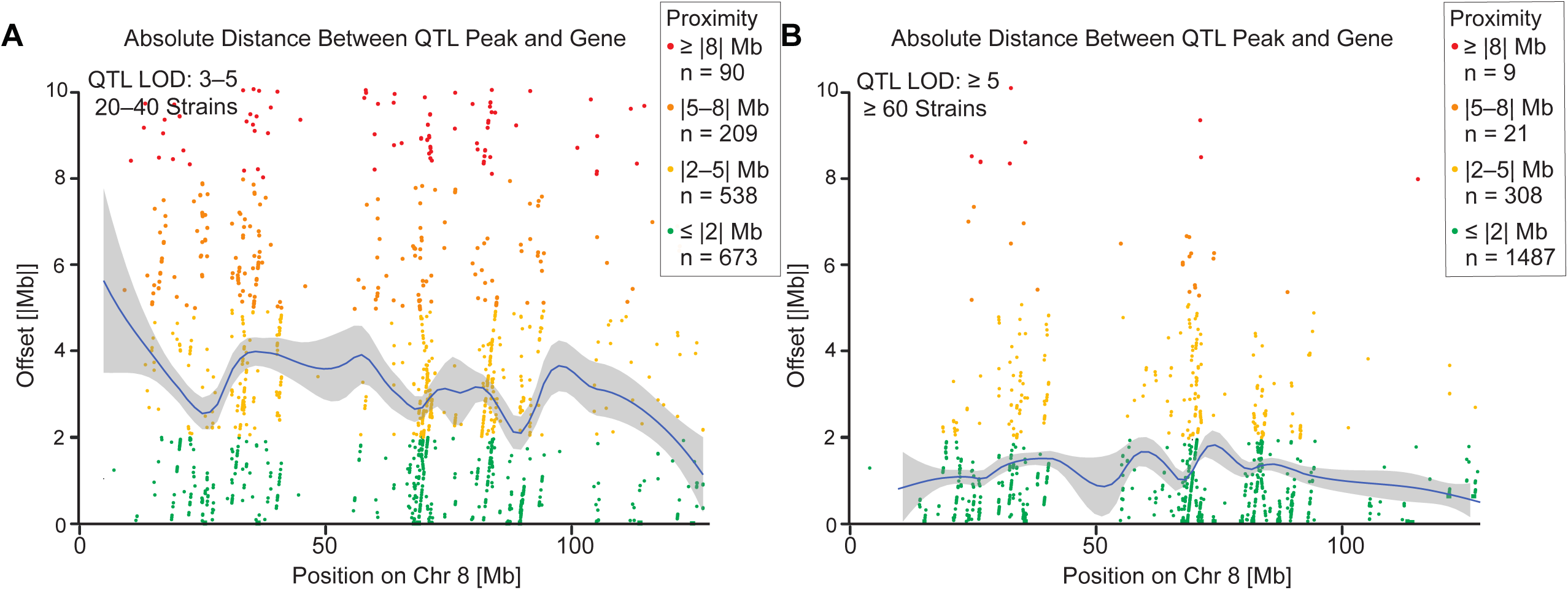
Empirical QTL mapping precision estimated using cis-acting expression QTLs. The distance between a probe and an eQTL is shown by each point. **(A)** Illustrates the precision achieved when mapping modest QTLs with LOD scores between 3 and 5 and using only 20 to 40 strains and minimal replication (usually 2 to 4 samples per strain). The number of replicates within each strain will affect the precision to a greater extent for QTLs with low heritability and modest LOD scores by increasing the effective heritability (see Belknap 1998). eQTL studies generally use only 1–4 replicates, so these precision values are conservative. Mean offset across the genome is 4.67 Mb, median 2.72 Mb. **(B)** Illustrates the precision achieved when mapping highly significant QTL QTLs with LOD scores greater than 5, and using between 60 to 80 strains, with at least 2 replicates. Mean offset across the genome is 1.38 Mb, median 0.76 Mb.

The mean precision for 17,805 eQTLs with LOD scores above 10 is ±0.96 Mb using transcriptome datasets containing only 60–80 BXDs. The median precision is 0.52 Mb. In other words, the offset between the location of the gene (using the more proximal end as a reference point), and the location of the SNP with the highest LOD score across the entire genome is generally less than 1 Mb. There is naturally high regional variation in precision (Figure 4). Most of the intervals with low precision are associated with intervals in which we have not yet defined locations of all recombinations, a problem that is now being solved by new sequence-based infinite marker maps ^55^. There may also be problematic regions of BXD genomes that contain large structural variatants (duplication, inversions, etc) compared to the reference genome.

It is possible to achieve subcentimorgan mapping precision with the BXD family, using only half of the full set of BXDs (Figure 4). Once genotypes are perfected by full genome sequencing we expect that with 60–100 strains users will achieve a mean resolution of 500 Kb and a median resolution of 250 Kb for Mendelian loci. Three factors contribute to this surprising precision: 1. the well balanced distribution of parental alleles across the genome (minor allele frequencies close to 0.5); 2. the ability to boost the effective trait heritability by resampling ^47^; and 3. the comparatively high density of recombinations archived in 150 BXDs thanks in part to the use of AI-derived strains (Figure 2). This level of precision does not differ appreciably from that typical of the Collaborative Cross, the Diversity Outbred, or even highly recombinant HS stock. In any case, precision much finer than this will often not be critical. The fuzzy functional boundaries of genes and the often high density of variants in linkage with each other shifts the burden of proof from pure mapping strategies to functional genomics, comparative analysis of human cohorts, complementary animal models, and to direct pharmacological and genetic engineering ^56^.

### The power of mapping with the BXD family

An analysis of statistical power is useful to estimate numbers of replicates and strains needed to detect and resolve major sources of trait variance and covariance. To start with the conclusion—it is almost always better to study small numbers of as many strains as can be obtained given the logistics of breeding, transportation, and experimental design ^2, 47^. Studying as few as two to four of each of 100 or more strains may strike many as counterintuitive but this is the right approach even for traits with low heritability. This fact has been the single most important motivation for increasing the number of BXD strains ^2, 47^.

In the context of mapping traits, an effective sample size can be estimated using an equation developed in Belknap and colleagues (1996). A more versatile method has been developed by Sen and colleagues ^57^ and implemented in the R package *qtlDesign* based on the H^2^_RIx̅_ ^47^. This heritability of strain means, in contrast to the heritability for individuals, is defined as H^2^_RIx̅_ = V_a_/(V_a_ + V_e_/n), where *V_a_* is the genetic variability (variability between strains), *V_e_* is the environmental variability (variability within strains), and *n* is the number of within line replicates. This equation highlights the usefulness of isogenic cohorts such as the BXD: the same genometype can be sampled multiple times, increasing the effective heritability. We rename this H^2^_RIx̅_ to H^2^_ix̅_ because it applies to all isogenic strains, not just recombinant inbred strains, for example diallel crosses (DXs, also known as RI intercrosses; RIXs) ^58, 59^ or hybrid diversity panels of inbred strains.

Given an estimated heritability and effect size, what is the optimum number of strains and number of biological replicates to use, given the need to maximize power and minimize the number of animals being used? To answer this, we have developed a power app to allow the user to examine these interactions in a two-founder recombinant inbred population, such as the BXD (https://dashbrook1.shinyapps.io/bxd_power_calculator_app/). This tool provides a clear visual demonstration that when heritability is low to intermediate, the gain in power from increasing the number of replicates is significant up to about four replicates and then tails off rapidly when heritability is higher than 0.5. Unsurprisingly, at heritabilities greater than 0.5 power is high even with as few as two replicates (Figure 5).

**Figure 5:**
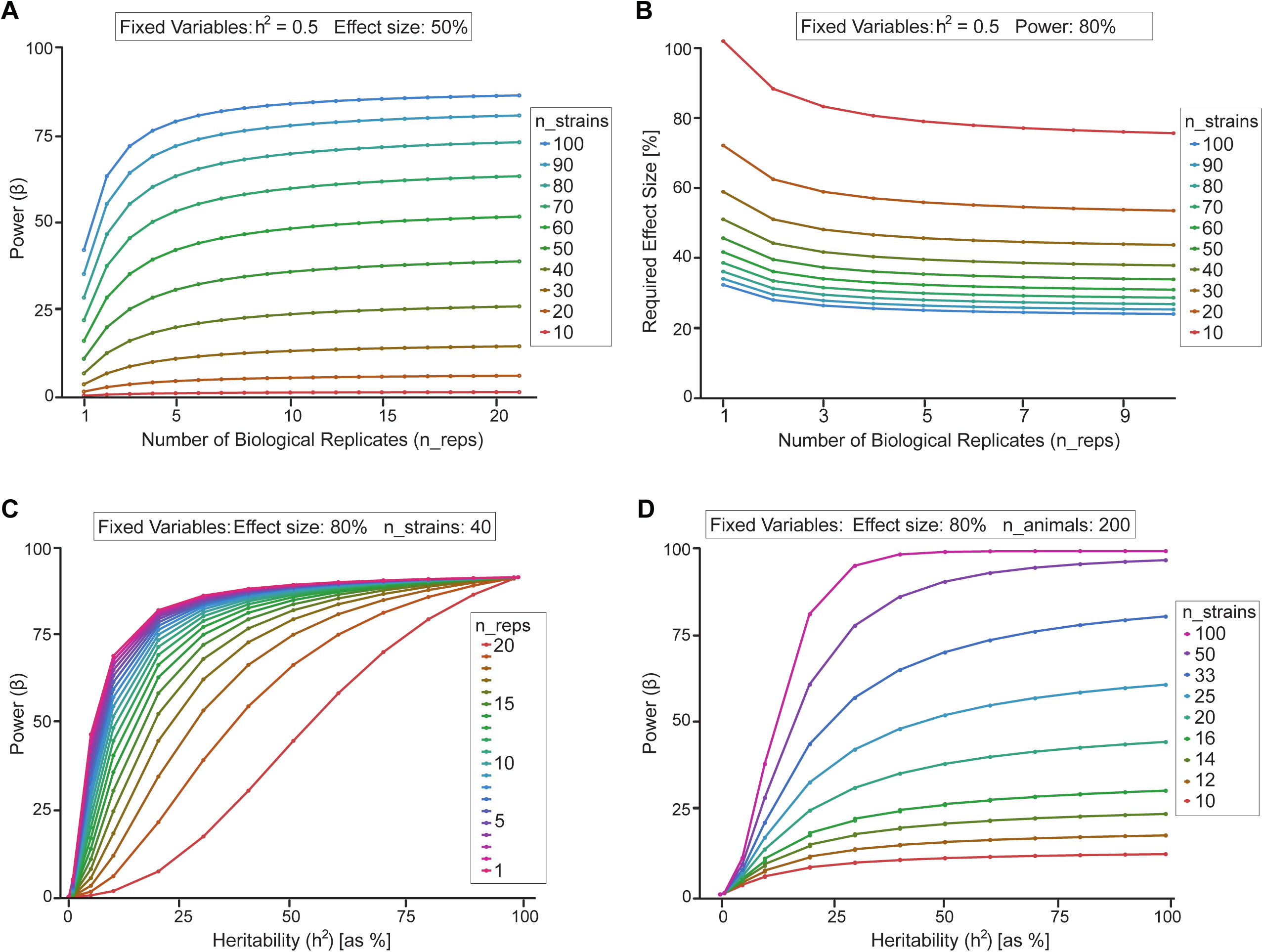
The relations between power, number of strains and within-strain replicates in recombinant inbred strains generated using qtlDesign (Sen et al., 2007). **A:** Power to detect a given effect for different numbers of replicates and different numbers of strains. Allele effect size and heritability were kept constant at 0.5. This shows that more replicates increase power but beyond four replicates the gain in power is marginal. At very low effect sizes power is low anyway, and then increases for intermediate levels. However, with large effect sizes, beyond 2 replicates very little power is gained. **B:** Effect size detectable, given different numbers of replicates and strains. Heritability was kept at 0.5, power was kept at 0.8. **C:** Power to detect a given effect size, dependent upon heritability and number of biological replicates. Allele effect size was kept at 0.8; number of BXD strains was kept at 40. The figure illustrates that with very low *H^2^* there is no power, and that with lower *H^2^* the gain in power is stronger with increasing number of replicates compared with high *H^2^*; again, beyond four replicates, the gain in power is marginal. **D:** Power to detect a given effect size, dependent upon heritability, with a total of 200 animals. These range from 200 lines with 1 replicate to 10 lines with 20 replicates. The figure shows that in contrast to what one might assume, even for low levels of *H^2^*, power always increases with more lines rather than more replicates. Effect size was kept at 0.8.

Investigators have to decide on the most effective use of limited resources, and this means decisions have to be made regarding the optimal design in terms of numbers of replicates and number of lines ^5^. For example, when limited to phenotyping 200 animals, the investigator must choose between phenotyping 10 lines with 20 replicates or phenotyping 40 lines with 5 replicates (reviewed in Williams and Williams, 2017). The figure and table in the power app reveals that for all levels of heritability it is always better to maximize the number of genomes rather than the number of replicates. Thus, even when heritability is low, increasing the number of replicates when more lines are available will not increase power to the same degree as increasing the number of lines. However, phenotyping more replicates will yield more reliable within-genotype variation estimates. Therefore, when using the expanded BXD panel we recommend phenotyping three to six replicates per line, when possible, using a balanced number of males and females. This is also optimal from a logistical standpoint, since it is often easier to obtain small numbers of animals for many strains from the Jackson Laboratory and other breeding centers. There is one exception to this advice. If one intends to select extreme response or outlier genomes for detailed analysis, then it is advisable to perform more extensive replication of these exceptionally interesting strains. In this context it should be noted that in samples with a small number of strains, the effect sizes of loci tend to be severely overestimated ^60, 61^. The utility and efficiency of RI and DX cohorts relative to F2 and heterogeneous stock cohorts is improved when heritability is low and numbers of replicates within strain are minimized (see Figure 3 from ^47^). Other factors to consider are genotyping costs, the resolution of genetic maps, and easy access to phenome data.

## Discussion

### Kinship relations and genetic drift

The expanded family of BXDs is a powerful resource for both forward and reverse genetic analysis of genome-to-phenome linkage. As this family has grown in size, relations among individual strains have become complex. Mapping algorithms that do not account for kinship (e.g. Haley-Knott methods) are no longer appropriate. We find it almost essential to use linear mixed models ^62–64^ or nonparametric equivalents such a mixed random forests ^65^ that account for kinship, epoch, and other cofactors. Several of these methods are now accessible in GeneNetwork 2 ^46^ and R/qtl2 ^66^.

The BXDs have kinship at several levels. First, there is random fixation and selection during the process of inbreeding ^23^. Second, there are notable differences in kinship among AI-derived strains, including BXD43 to BXD102 and BXD160 to BXD186 ^20^. This kinship has its origins mainly in the 8–14 generations of intercrossing that preceded inbreeding ^22^. Third, there are substrain groups highlighted by common numbers but unique substrain codes (e.g. BXD24/TyJ, BXD24/Tyj-*Cep290*<*rd16*>/J, BXD29/TyJ and BXD29-*Tlr4*<*lps*-2J>/J). Both pairs are co-isogenic, and carry unique spontaneous mutations. In contrast, the set closely related BXD48s; BXD65s, and BXD73s resulted from production errors. Fourth, there is kinship based on epochs. Each epoch of BXDs were generated with unique B6 and D2 parents, albeit nominally the same strains. These parents, but more importantly their progeny BXD strains, differ due to fixation of spontaneous mutations inherited from both parents. Finally, each strain will fix a variable number of rare spontaneous mutations during inbreeding. While these rare fixed mutations do not produce an epoch effect, they can produce remarkable outlier strains.

Shifman and colleagues ^67^ detected a surprisingly large number of new variants (*n* = 47 out of about 13,000 SNPs) in the set of strains generated by Taylor in the early 1990s (BXD33 though BXD42), and a small number (*n* = 5) of even more recent mutations in BXDs generated in the late 1990s at UTHSC. While the majority of mutations are neutral or non-functional, we already know that a handful produce interesting differences in gene expression and higher order phenotypes among different phases of the BXD family ^68, 69^. Analysis of expression data from different epochs highlights ∼50 genes modulated by strong cis-acting expression QTLs only in newer strains. For example, introduction of a premature stop codon in glycoprotein nmb (*Gpnmb)* and deletion of the last coding exon in killer cell lectin like receptor D1 (*Klrd1*; CD94) occurred spontaneously in the DBA/2J paternal strain in the 1980s ^70, 71^. The result is a loss of function through truncation of each protein and both genes are associated with strong cis eQTLs, but only in the newer strains. The same phenomenon is true of spontaneous mutations that occurred in the 1980s in the C57BL/6J maternal parent that led to the deletion of *Nnt* exons and a striking reduction in *Gabra2* expression in brain ^68, 72, 73^.

### Improved mapping of BXD phenome and omics data

The BXD were first used to map trait variants segregating in the parental strains in the early 1970s. Even with sparse genotypes, Taylor and colleagues were able to map a cadmium toxity locus to a 24 cM interval ^1^. We compared the pre-2001 ‘original’, 2001-2016 ‘classic’, and post-2017 ‘current’ genotype files. The latest file increases the number of QTLs detected at a LRS threshold of 20 by ∼25%. As a specific example, using the latest genotypes and the GEMMA mapping algorithm, Taylor’s 1970s data (BXD Phenotype 13035) now gives a map with a 2-LOD confidence interval of 3 Mb. When this map is combined with SNP data for 29 common strains that Taylor and colleagues also generated in their landmark study (GN 49785 in the Mouse Diversity Panel cohort) this locus can be restricted to a 400 Kb region centered at SNP rs13477430 that includes *Slc39a8*, a heavy metal transporter expressed almost exclusively in the testes. This locus was formally cloned by Dalton and colleagues in 2005, although the causal variant has still not been defined functionally ^74^.

This highlights two important points. First, it is possible to map Mendelian traits and even quantitative traits with modest LOD scores with good precision, even when using a small numbers of strains ^75–77^. Second, a good way to transition from QTLs to specific genes, variants, and mechanisms is often to use complementary resources such as panels of common inbred strains, Collaborative Cross (CC), or Diversity Outbred (DO) cases, efficient screens of candidate genes using *in vitro* and *in vivo* assays ^48, 76^, and even human genome-wide association study (GWAS) data ^78–82^. But the most obvious way to improve precision and power is to simply use a larger numbers of strains.

Cloning QTLs and quantitative trait (QT) nucleotides is becoming significantly easier thanks to this excellent precision, improved genotypes ^23, 67, 83, 84^, full genome sequence for both parental strains ^3, 43^, powerful omics resources for the BXDs ^4, 27, 85, 86^, better mapping algorithms that account for cofactors and admixture ^62, 87, 88^, and far more efficient molecular validation methods ^45^. The BXDs have been used to clone roughly 20 QTLs, including two tightly linked genes, *Iigp2* and *Irgb10,* for *Chlamydia* infectivity ^89, 90^, *Fmn2* as a master controller of tRNA synthetases in neurons ^91^, *Ubp1* for blood pressure ^78^, *Hc* for H5N1 influenza resistance ^92^, *Comt* as a master controller of neuropharmacological traits ^93^, *Alpl* for hypophosphatasia ^2^, *Mrps5* for longevity ^76^, *Bckdhb* for maple syrup urine disease, *Dhtkd1* for diabetes ^27^, *Hp1bp3* for cognitive aging ^94^, and *Ahr* for locomotor activity ^95^.

### The BXD phenome is unrivalled for experimental precision medicine

The BXDs are currently the largest and most deeply phenotyped vertebrate genetic reference panel, and have been used by well over 200 groups worldwide. Researchers benefit from access to an expanded BXD family, which has been increased by 88 strains to a total of 150 extant strains ^41^. Almost all of the data are freely accessible at GeneNetwork (https://gn2.genenetwork.org/) ^46, 87, 96^. The assembly of phenome data and their integration into a web service was started in 2001 ^21, 97^ and is still a core activity of our groups ^3^. This wealth of legacy data accumulated over the past four decades is represented by well over 1,000 publications, over 7,000 open phenotypic measurements, 150 open genome-wide molecular profiles that include proteome data sets generated under different metabolic conditions, and over three million mRNA measurements. The BXD transcriptome has been densely sampled across brain and peripheral tissue and is represented by more than 80 independent data sets using a wide variety of platforms— ^98, 99^. Recent work by multiple investigators includes the generation of miRNA, proteomic, metabolomic, epigenomic, and metagenomic data for the BXDs ^4^. This assemblage of quantitative data has been integrated into GeneNetwork and the Mouse Phenome database ^100, 101^ and is freely accessible both as individual data sets or in its entirety.

The size and depth of the multiscalar phenome constructed for the BXD family makes it an excellent resource for systems genetic and genomics, as well as a foundation for experimental precision medicine. The genetic architecture of one or many traits can be dissected and relations between traits exposed. The ability to resample genotypes across a stable genetic reference population enables the expansion of the phenome in almost any direction and including cellular and molecular phenotypes at any stage. There are two main obstacles: The first is the development of structured and controlled vocabulary or ontologies for phenotypes that allows for cross-species translation and bioinformatic analyses. The second problem sounds trivial but is actually difficult—getting a large and diverse community of biomedical researchers engaged in using replicable genetic resources and then in providing their extremely valuable core phenotype data and metadata to curators and data integrators.

The BXD phenome encompases multiple levels of data, from single molecules to complex behavioral repertoires, measured under standard or various environmental exposures, including alcohol and drugs of abuse ^31, 102–104^, infectious agents ^92, 105, 106^, dietary modifications ^81, 82, 107–110^, stress ^111, 112^ and as a function of age. While this represents the largest phenome for any group of animals it is still an unbalanced phenome and has significant gaps that need to be filled, for example, in embryology, cellular and systems physiology, cardiovascular and urogenital systems. Major areas of research in which the BXDs have experimental advantages over other systems are particularly in epigenetics and gene-environment interactions ^44^.

### Future Directions: Pleiotopy, epigenetics, gene by environment interactions and precision medicine in the extended BXD family

Most of the work carried out in the BXD family to date has had the simple goal of defining single causal gene variants and mechanisms that contribute to heritable differences in disease risk. But we need to tackle a complementary and far more challenging problem—the complex interplay among sets of variants, constellations of phenotypes (pleiotropy), and different treatments and environments ^4, 33, 48, 113^. Experimental cohorts such as the BXD family are ideal to achieve this goal, and are powerful experimental companions to human genetics and clinical research. These unique types of immortal murine families will make it possible to experimentally test the promise and limits of precision medicine.

Over the next decades large cohorts of humans and other species will be sequenced and analyzed using advanced molecular methods (e.g. ^114^). We will have access to sequence and electronic health records for millions of subjects (e.g. Million Veterans Projects ^115^ and UK Biobank; www.ukbiobank.ac.uk ^116, 117^), but will this be enough to address the main goals of precision health care—to accurately predict risk of disease before onset and to optimize prevention and therapeutics? Only when combined with advanced experimental systems such as the BXD panel or the Collaborative Cross will we be able to harness the full potential of conjoint multi-systems approaches ^118^.

Standard inbred strains of mice ^119, 120^ and their RI progeny such as the BXDs and the Collaborative Cross ^23, 59^ are a good resource with which to test the limits of precision medicine. They enable a community of researchers in disparate fields to gradually assemble coherent phenomes of great size for isogenic lines with nearly complete sequence data ^121^. The BXDs excel in this dimension. But BXDs and other murine populations should not be used to the exclusion of other resources. For example, BXDs and other isogenic lines of mice, including common inbred strains, the Collaborative Cross, and panels of congenic lines can all be used jointly. The statistical analysis of complex admixed cohorts is now computationally tractable ^122^ using the same algorithms used to handle admixed human GWAS cohorts. Williams and colleagues and Lusis and colleagues have been systematically using a “hodge-podge” of up to 100 isogenic strains (what is now called a hybrid mouse diversity panel) in joint systems genetic studies of many clinically relevant traits ^75, 123–127^. Indeed, combined mouse QTLs and human GWASs have identified novel candidates and pathways illustrating the translational value of experimental data from RI populations ^2, 77, 78–82, 128–130, 131^.

The BXD family of 150 strains is now an appropriate number of genomes to evaluate predictive modeling of biological processes and disease risk, and for developing computational models of genome-to-phenome relations ^3, 90, 132^. In a two-parent RI family such as the BXDs minor allele frequency is of course close to 0.5 for most variants. Furthermore, there are only four two-locus combinations (BB/BB; BB/DD DD/BB, and DD/DD). Thus with 100 strains each of these four combinations will typically (95% of the time) be represented by 16 or more strains. The BXDs are therefore large enough to enable the analysis of epistatic interactions with good power. But the power to study epistasis can be greatly amplified by crossing BXD strains to generate a DX. With a subset of 100 BXD strains, it is possible to generate 9900 reciprocal pairs of DX F1 projeny. For example, crosses of female BXD99 to male BXD100 and vice versus give rise to 99X100 and 100X99 isogenic DX projeny. These reciprocal DX pairs generally do not differ for either mitochondrial or Chr Y since these are both fixed as *B* and *D* haplotypes, respectively, in the BXDs (but there are exceptions).

Reciprocal diallel crosses (reciprocal DXs) have been used to investigate parent-of-origin effects on ultrasonic vocalization ^14^. In F1 reciprocal crosses between most isogenic strains, females have identical nuclear genomes, differing only in their mitochondrial genome, whereas males will also differ in their sex chromosomes. This allows the investigation of parent-of-origin and sex chromosome effects ^133^. This makes them almost uniquely suited for the investigation of parent-of-origin effects. As well as expanding the number of isogenic mice in the BXD family from 150 to over 22,000, BXD animals can also be crossed to other strains. Isogenic strains can be crossed to traditional mutant mouse models to investigate the effect of diverse genetic backgrounds on mutations linked to human diseases ^134–136^. Neuner et al. crossed a subset of the BXD panel to the transgenic and humanized Alzheimer’s Disease (AD) 5XFAD model ^137^ to produce F1 hybrid transgenic AD-BXDs, and were able to demonstrate a profound effect of genetic background on both cognitive and physiological phenotypes ^134^. This provides an entirely new avenue of investigation using resources such as the animals generated from the International Mouse Phenotyping Consortium (IMPC; http://www.mousephenotype.org/) ^138^, Mutant Mouse Resource & Research Centers (MMRRC; https://www.mmrrc.org/) and Knockout Mouse project (KOMP; https://www.komp.org). By crossing humanized or interesting genetically engineered lines on a single genetic background to genetically defined individuals from a diverse population, variants which modify traits of interest can be mapped, thus elucidating epistatic interactions, and providing information on disease course, not just disease risk. Again, this provides a model that approximates a human population, with the exception that individual genomes can be resampled infinitely.

We now need to rethink the best uses of mouse models for complex trait analysis. The main reason is that the original challenge of achieving high mapping precision has been largely overcome in human genome-wide association studies (GWAS Catalog at www.ebi.ac.uk/gwas). Tens of thousands of variants have been mapped to small haplotype blocks—usually under 200 Kb—for a broad spectrum of human traits—from differences in mRNA expression to stature and schizophrenia ^139–141^. This is only the beginning. The precision and the power in human genetics will improve greatly over the next several decades as full genome sequences, better human disease phenotyping, and electronic health records are merged at the scale of millions of subjects and whole nations. Therefore, we need to revamp experimental genetic resources in an era flooded in GWAS hits. How are new and old mouse resources best repositioned to help deliver on the still unmet and much more integrative promises of predictive genetics and personalized precision health care?

The short answer is that we need large genetically complex resources with matched multiscalar and multisystems phenome data that can be used to test and predict effects of interventions, drugs, and other gene-by-environment interactions. We need resources that define and test progressively more sophisticated computational models as a function of genotype, case history and exposure. We need replicable mouse populations with the same level of genetic complexity as human cohorts but without the many disadvantages of clinical research—high cost, poor compliance and control, confidentiality constraints, and importantly, ethical limitations on experimental procedures. Mouse populations—including large families of recombinant inbred strains and their isogenic but non-inbred RI F1 diallele crosses (DX cohorts)—are ideally positioned to be the experimental test bed with which to define the power and limits of precision health care.

### Conclusions

Systems genetics using mouse models has been revitalized over the last decade thanks to several new resources, including the BXD family of recombinant inbred strains ^20^, the Hybrid Mouse Diversity Panel (HMDP; ^121, 126^, and the Collaborative Cross (CC) ^142–145^. Mouse models stand as an excellent resource to test hypotheses and explore mechanisms in the new era of precision medicine, working in combination with new genomics tools available in human cohorts. The main limitation has been relatively modest mapping power and precision—a simple problem caused by small numbers of strains. With 150 strains now readily available, the BXD family has overcome this problem.

## Methods and Materials

### BXD strains

Between 2009 and 2010 we initiated 74 BXD strains (BXD104 to BXD186). We initiated another 34 strains in 2013 (BXD187 to BXD220). We used both conventional F2 intercrosses (*n* = 88) and AI progeny (*n* = 20) to make the 108 new lines. BXD160 through BXD186 were derived from unique matings of AI progeny gifted us by Dr. Abraham Palmer at G8 and G9 late in 2010 (Figure 1, Supplementary figure 1, Supplementary table 1).

In cases of low reproductive fittness, we often attempted to rescue lines by outcrossing young male BXDs to C57BL/6J females followed by three of more sequential backcrosses (N3) to the RI male to produce progeny enriched for the BXD genome. In more recent cases, pairs of at risk strains were crossed to produce RIX progeny ^23, 50^, which were then inbred by sibling mating. BXD221 to BXD228 are RIX-derived strains of this type (Supplementary figure 1, Supplementary table 1).

Unless otherwise specified, all animals used in this study were raised in a closed-barrier pathogen-free vivarium at the University of Tennessee Health Science Center. All BXD strains are available under a standard material transfer agreement; the most important limitation being that they cannot be sold or distributed without approval of the Jackson Laboratory or UTHSC. Availability information on all BXD lines can be found in Supplemental Table 1.

### Genetic map construction and genotype error correction

Individuals from each of the 198 BXD lines were genotyped in late 2011 and in late 2015 using the Neogen/GeneSeek MUGA or GigaMUGA arrays. These were combined with previous genotypes generated using Affymetrix and MUGA platforms. Unknown genotypes were imputed as B (C57**B**L/6J-like) or D (**D**BA/2J-like), or were called as H (heterozygous) if the genotype was uncertain.

The full dataset contains 37,000 markers and was optimized for mapping efficiency by excluding markers showing identical strain distribution patterns. Markers that flank sites of recombination were retained. As a result, strain distribution patterns are often defined by proximal-distal marker pairs. Whenever possible, we verified and updated genotypes of the original BXD strains (BXD1 - BXD102) to reflect those of stock available from the Jackson Laboratory (^83^ and from Petr Simecek and Gary Churchill; http://cgd.jax.org/datasets/diversityarray.shtml).

Genotypes for all strains were smoothed and curated to remove highly implausible (double) recombination events, e.g. unsupported genotypes (singletons) that introduced two recombination events over less than 100 kb ^23^. In general, we imputed unknown or heterozygous genotypes on the basis of flanking markers. Undefined genotypes between recombinations were coded as heterozygous, and telomere genotypes were imputed using the closest flanking marker.

In the newer BXD strains there are many regions that are still heterozygous. Generated genotypes using standard platforms still show regions of low marker density and a high frequency of recombinations. These regions of low marker density were filled with imputed genotypes using *cis*-eQTLs of genes in the problematic intervals. Microsatellites and *cis*-eQTL genotypes were generated by the Williams/Lu laboratory.

The assembled and error checked genotype file includes 7,324 markers for 191 independent strains, and 7 substrains, has been available since January 2017 at http://www.genenetwork.org/genotypes/BXD.geno and is the default genetic map used when QTL mapping on GeneNetwork (http://www.genenetwork.org).

### Computing centi-Morgan positions

For each chromosome we compute the number of observed recombinations from one position to the next, ignoring the heterozygous and unknown genotypes (nRecP). We then compute the recombination fraction (R) by dividing the number of recombinations observed (nRecP) by the population size used to generate the genetic map. The recombination fractions (R) are then deflated using the formula:

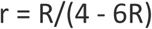

which is the inverse of the formula R = 4r/(1+6r) for a recombinant inbred line by sibling mating. We then use the function imf.cf available in the R package R/qtl ^64, 146^ to convert from centiMorgan using the Carter-Falconer map function^147^. Positions obtained start at 0 on each chromosome, requiring all positions to be shifted so that the first marker on the map matches the known centiMorgan position of the first marker on the genetic map. The old genetic map and the mousemapconverter (http://cgd.jax.org/mousemapconverter/) were used to obtain centiMorgan positions for the first marker on a chromosome, all position on a chromosome are shifted to align the genetic map to the known starting positions of the markers. Code for computing marker positions is available in the BXDtools package (https://github.com/DannyArends/BXDtools), and positions can be recomputed using different map functions when required (e.g. Haldane ^148^, Kosambi ^149^, Morgan ^150^).

### Genome wide visualization of recombinations

For each chromosome we compute the number of observed recombinations per strain from the start of the chromosome (nRecS), ignoring the heterozygous and unknown genotypes. We then visualize this by plotting the number of recombinations for each BXD strain per chromosome using a color gradient. Strain numbers are used for the ordering on the X-axis, and horizontal lines are added to visually separate the different epochs and annotate the epochs on the X-axis.

### Comparing genotype files

During the BXD’s 40+ years of existence, several genotype files have been used for QTL mapping. The first, genome-wide map (which we call the ‘original’) was used pre-2001, and contained 1578 microsatallites ^23^. The second ‘classic’ map was used between 2001-2016 and contained 3811 SNPs from various microarrays ^151^. The third, ‘current’ post-2017, used 7321 SNPs and is described above.

For the pre-2001 microsattalites, Mouse Genome Informatics was used to determine updated (mm10) positions for 1419 of the probes, but this left 159 without mm10 postions. We used NCBI Probe database (https://www.ncbi.nlm.nih.gov/probe/) to search for these 159 probes, and for 107 we were able to get primer sequences. For these sequences, we than ran Primer-BLAST (https://www.ncbi.nlm.nih.gov/tools/primer-blast/index.cgi) to determine where in the mm10 genome they would amplify. For the remaining 52 markers, we used BLAST to align the target sequence to mm10 to identify the mm10 position. This left 11 markers which Primer-BLAST either predicted a different chromosome would be amplified, or which did not have any mm10 genomic targets.

These ‘original’ genotypes only exist for the first 36 BXD strains (BXD1-42), and so the remaining strains were imputed (BXD24a and BXD 43+), using the genotype at the microsatalite’s position in the ‘current’ post-2017 genotyping file. To determine our power to detect QTLs, we ran mapping using QTLreaper for the 7562 phenotypes collected for the BXD, and the number of phenotypes which had an LRS greater than three threshold values: 15, 20 and 25. These correspond to suggestive, significant and highly significant LRS values. The larger the number of phenotypes above these thresholds, the greater our power to detect QTLs.

### Precision and Resolution

To estimate empirical precision of mapping across the genome we have extracted data from 40 large-scale genetic studies of gene expression of the BXD family. These studies used between 21 and 79 strains with variable numbers of within-strain replicates, over 27 tissues. We defined eQTLs that map within ±20 Mb of the gene associated with the transcript’s measurement as a *cis*-acting expression QTLs (*cis*-eQTLs). Unlike a standard F2 intercross, this ±20 Mb window is roughly equivalent to a recombination distance of ±40 cM in the highly recombinant BXD family, and the statistical association between markers this far apart is generally quite low (*r*^2^ less than 0.3).

Precision was estimated using the offset between the location of the gene (using the 5’ end of the probe or probe set as a reference point), and the location of the SNP with the highest LOD score. Again, this is conservative, as the causal variant is unlikely to be exactly at the 5’ end of the gene. There are often two or more neighboring SNPs with equally high LOD scores and we simply take the most proximal marker to compute offset. The light blue local regression (LOESS) smoother curve ^152^ was computed using a window with a size of 0.333% of the genome. Smoothing was carried out and data was plotted using the *ggplot2* R package.

### QTL mapping

QTL mapping using GeneNetwork has been described in detail elsewhere ^87^. However, in brief, quantitative trait loci (QTLs) are segments of the genome affecting a particular phenotype ^153^. QTL mapping, identifying QTLs to explain the genetic basis of complex traits, relies on being able to make correlations between genetic markers and phenotypic traits in a population. Individuals are scored for their phenotype for a particular trait, and their genotype at a marker. If there is a difference in mean phenotype between those individuals with one genotype compared with the other than we can infer that there is a QTL linked to that marker. If there is no difference between the means we can conclude that the loci does not influence the phenotype in that population ^153, 154^.

Due to the very high density of markers, the mapping algorithm used to map BXD data sets has been modified and is a mixture of simple marker regression, linear interpolation, and standard Haley-Knott interval mapping ^155^. When two adjacent markers have identical strain distribution patterns, they will have identical linkage statistics, as will the entire interval between these two markers (assuming complete and error-free haplotype data for all strains). On a physical map the LRS and the additive effect values will therefore be constant over this physical interval. Between neighboring markers that have different strain distribution patterns and that are separated by 1 cM or more, we use a conventional interval mapping method (Haley-Knott) combined with a Haldane estimate of genetic distance. When the interval is less than 1 cM, we simply interpolate linearly between markers based on a physical scale between those markers. The result of this mixture mapping algorithm is a linkage map of a trait that has an unusual profile that is particular striking on a physical (Mb) scale, with many plateaus, abrupt linear transitions between plateaus, and a few regions with the standard graceful curves typical of interval maps.

Prior to the 2017 release of the genotypes described in this manuscript interval mapping in GeneNetwork relied on 3,795 informative SNP markers across all autosomes and the X chromosome ^151^. These markers were generated using the MUGA array in 2011, along with earlier generated genotypes on the Affymetrix and Illumina platforms ^67^. As described above, loci are identified in GeneNetwork by the computation of a likelihood ratio statistic score and significance was determined using at least 5,000 permutations of the phenotype data.

Updated QTL mapping methods, such as R/qtl2 ^66, 146^, Multiple QTL mapping ^64^, GEMMA ^156^ and pyLMM ^63^, have been implimented on the GeneNetwork2 site ^46^. GEMMA, and pyLMM allow to account for kinship between BXD strains by computing a genetic kinship matrix given specific strains used in an analysis, allowing for precise kinship correction and improved QTL mapping performance when mapping any of the BXD phenotypes in the GeneNetwork database.

### Phenome wide association (PheWAS)

Phenome-wide association studies (PheWAS) associates a single genetic marker with a rich set of phenotype data, to investigate a phenotypic and pleiotropic effects of a genetic marker. PheWAS is related to GWAS, and PheWAS can be characterized a GWAS on its side, investigating if the genetic variants linked to this marker influence multiple phenome categories. Phenotype data is first grouped in so called phenomes, groups of phenotypes which are functionally, physiologically or genetically related to each other. To visualize the PheWAS scan, phenotypes are put on the X-axis, and strength of association (either in LOD or LRS) is put on the Y-axis. Significance thresholds were computed and adjusted based on Benjamini-Hochberg correction ^157^, where the number of tests is equal to the number of phenotypes in the PheWAS scan. The term PheWAS was first coined by Arends et al. while investigating longitudinal electronic medical records (EMRs) coupled to genetic data ^158^. The (extended) BXD population is the most densely phenotyped and manually annotated inbred mouse population, and the extended BXD population allows for PheWAS with unprecedented precision. PheWAS is not yet integrated in the original GeneNetwork, but is available in the BXDtools package (https://github.com/DannyArends/BXDtools) and on the Systems-Genetics.org website (https://www.systems-genetics.org/) which will be integrated into GeneNetwork2 allowing for point and click PheWAS scans of the BXD phenome containing ∼ 7000 classical phenotypes. Systems-Genetics.org also contain the tools for Mediation, reverse mediation and gene expression PheWAS (ePheWAS).

### Power calculation

Power calculations were carried out based on the method of Sen and colleagues ^57^ and implemented in the R program *qtlDesign* based on the H^2^_RIX_ defined by Belknap ^47^, which we renamed H^2^_ix̅_ as it is applicable to any isogenic strain, not just recombinant inbreds. The *Detectable* function was used, to determine the power available with an RI design to detect a locus of a given effect size.

For calculation of the H^2^_ix̅_ four variables need to be known: the broad sense heritability (H^2^), the number of within-strain replicates (biological replicates), the number of strains used and the locus effect size (the proportion of total genetic variance explained by the locus). To calculate H^2^, the genomic variance (gen.var) was kept constant at 1, and the environmental variance (env.var) was varied to produce H^2^ values ranging between 0 and 1. The number of biological replicates and number of strains are self-explanatory, and were capped at 10 and 150 respectively, since >10 replicates produced a marginal increase in power, and because there are 150 BXD lines currently available.

The locus effect size is the amount of the variance which is due to genetics (gen.var) explained by a single QTL. That is, a value of 0.2 would mean that a QTL explains 20% of the genetic variance, and a value of 1.0 indicates a Mendelian locus (one QTL which explains all of the genetic variance).

The power given is the ability of the experiment to correctly detect a true positive QTL, given the other values above. The BXD power app can be found at https://dashbrook1.shinyapps.io/bxd_power_calculator_app/ and uses the *Shiny* R package (https://CRAN.R-project.org/package=shiny)

## List of abbreviations

AD: Alzheimer’s Disease
AI: advanced intercross
B6: C57BL/6J mouse strain
BLAST: Basic Local Alignment Search Tool
BXD: BXD family of recombinant inbred strains of mice, produced by crossing the B6 and D2 parental strains
CC: Collaborative Cross
Chr: Chromosome
cM: centimorgan
D2: DBA/2J mouse strain
DO: Diversity Outbred
DX: diallel cross
EMRs: electronic medical records
eQTL: Expression quantitative trait locus
F2: Filial generation 2
GEMMA: Genome-wide Efficient Mixed Model Association
GigaMUGA: Giga Mouse Universal Genotyping Array
GN: GeneNetwork (http://genenetwork.org)
GN2: GeneNetwork2 (http://gn2.genenetwork.org)
GWAS: genome-wide association study
H^2^_ix̅_: heritability of isogenic strain means
H^2^_RIx̅_: heritability of recombinant inbred strain means
HMDP: Hybrid Mouse Diversity Panel
IMPC: International Mouse Phenotyping Consortium
JAX: The Jackson Laboratory
Kb: kilobase
KOMP: Knockout Mouse project
LOD: logarithm of odds
LRS: likelihood ratio statistic
Mb: Megabase
MMRRC: Mutant Mouse Resource & Research Centers
MUGA: Mouse Universal Genotyping Array
nRecP: number of observed recombinations from one position to the next
nRecS: number of observed recombinations per strain
PheWAS: phenome-wide association studies
QT: quantitative trait
QTL: quantitative trait locus
*r^2^*: Coefficient of determination
RI: Recombinant inbred
RIXs: Recombinant inbred intercrosses
SE: Standard error of the mean
SNP: single nucleotide polymorphism
UTHSC: University of Tennessee Health Science Center

## Declarations

### Availability of data and material

The datasets generated and/or analyzed during the current study are available in the GeneNetwork repository, https://www.genenetwork.org/

## Acknowledgements

Our thanks to Dr. Abraham Palmer for providing us with valuable B6D2 advanced intercross progeny (G8 and G9) in 2011 that we used to make BXD160 through BXD186. We thank Dr. Benjamin A. Taylor for initiating the BXD family and for his continued support and encouragement. We thank members of the EPFL Center for Phenogenomics in particular Drs. Xavier Warot and Emilie Gesina for sharing data from their BXD colony, and Erik Soehnel from Scionics for his assistance in obtaining the full breeding statistics. The UTHSC Center for Integrative and Translational Genomics (CITG) has supported the production of the BXD colony at UTHSC and will continue to support this colony for the duration of the grant. The CITG also provides generous support for computer hardware and programming associated with GeneNetwork, and our Galaxy and UCSC Genome Browser instances. Finally, the CITG as well as the UT-ORNL Governor’s Chair endowment to RWW will help support the sequence analysis of the BXDs. This work is required to extract the interesting but small subset of rare private alleles that each strain accumulates during inbreeding.

## Supplementary files

**Supplementary figure 1**

**.png**

**Production of each epoch of the BXD recombinant inbred family.**

The BXDs up to BXD220 have been produced in 6 epochs, with epoch 1 being started in 1971. Red coloring has been used to represent regions of the genome coming from the inbred C57BL/6J (B6) parental strain, whereas white coloring has been used to represent regions of the genome coming from the inbred DBA/2J (D2) strain. Solid lines have been used to represent a single generation, whereas dashed lines represent several generations, with the number of generations written along the line. During the 1970s to 2001s strains that were at risk of going extinct were often backcrossed to B6 in an attempt to rescue them, and this is shown in the figure. During epochs 4-6, strains at risk of extinction were sometimes crossed to other BXD RI strains in an attempt to rescue them.

Beneath each epoch, the number of inbred strains begun, the number of strains with data within GeneNetwork, and the number of strains extant are shown. In cases where any crosses or backcrosses occurred these are also noted. Although the majority of the BXD family have the D2 Y-chromosome and B6 mitochondrial genome, this is not the case for all strains, and this is noted as well.

Adapted from (Nadeau and Auwerx, 2019; Peirce et al., 2004; Williams and Auwerx, 2015). Full details can be found in Supplementary file 2.

**Supplementary table 1**

**.xlsx**

Information on the nomenclature, availability, derivation and genotyping of all BXD strains (BXD1-BXD228).

Shows alternative names that have been used for BXD family members, details of when breeding was started, when production of these strains began at the Jackson Laboratory (JAX), the stock number and availability from JAX, the epoch, method of derivation, the origin of the mitochondrial genome, and platforms that each strain has been genotyped on, and the generation number if known.

**Supplementary table 2**

**.xlsx**

List of annotations in the BXD phenotype ontology used for our PheWAS study. Shows the GeneNetwork trait ID, the ‘Phenosome’ (the numerical phenotype category), the phenotype category, the full phenotype description, the authors who produced the phenotype, the year of publication (or year of entry into GN if unpublished) and the PubMed ID (if published).

