## Supplementary figures and images for "The expanded BXD family of mice: A cohort for experimental systems genetics and precision medicine"

### Supplementary Figure 1

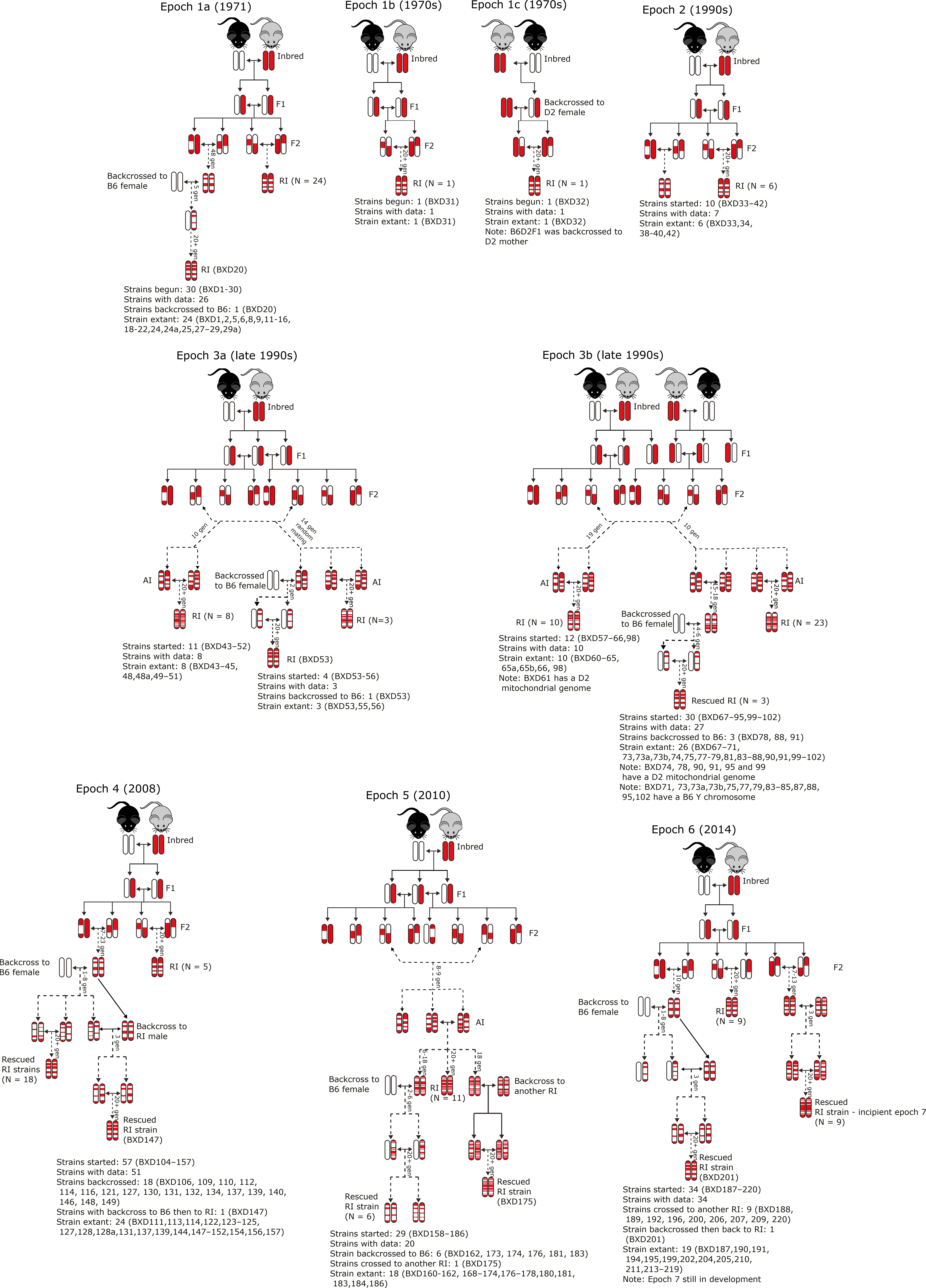
